# Motor cortex directly excites the output nucleus of the basal ganglia, the substantia nigra pars reticulata

**DOI:** 10.1101/2024.03.19.585632

**Authors:** William Scott Thompson, Sten Grillner, Gilad Silberberg

## Abstract

Inhibitory neurons of the substantia nigra pars reticulata (SNr) serve as a primary output through which the basal ganglia regulate behaviour. Projections to the SNr from beyond the basal ganglia have also been identified anatomically. Using a virally-targeted optogenetic approach, combined with whole cell patch-clamp recordings of SNr neurons in acute brain slices, we show that projection neurons of the primary and secondary motor cortices (M1 and M2) make functional excitatory synapses with subpopulations of inhibitory SNr neurons. Furthermore, we demonstrate that photostimulation of these cortical axon terminals increases SNr neuron firing rate. To further investigate the spatial organisation of cortical input to SNr, we employed a transsynaptic viral-labelling approach to identify SNr neurons receiving monosynaptic input from M1 and M2. We found a topographical relationship between motor cortex and SNr, and identified downstream targets of cortical-recipient SNr subpopulations. These findings reveal functional pathways by which M1 and M2 can directly modulate basal ganglia output to different downstream targets.

## Introduction

Successful motor output requires coordination between the plethora of brain structures that make up the motor system. The basal ganglia are a set of subcortical nuclei that integrate multiple streams of information, to select and reinforce context appropriate motor programs^1^. Located in the ventral midbrain, the substantia nigra pars reticulata (SNr) is the largest output nucleus of the rodent basal ganglia and serves as an important link between the integrative processing of the basal ganglia, and numerous motor executors in the midbrain and brainstem^2–4^.

GABAergic SNr neurons are intrinsically spontaneously active, and directly regulate behavioural output through tonic inhibition of their targets in the midbrain and brainstem^5–10^. In addition, they transmit efference copy information back upstream, via collaterals to thalamic nuclei^4,11^. Rather than forming a broad global output, distinct groups of GABAergic SNr neurons project to unique sets of targets^4^. These target-defined neuronal subpopulations are both spatially clustered and electrophysiologically distinguishable. SNr projection patterns reflect a modular organisation of the basal ganglia as a whole, whereby information flow is organised into parallel modules for specialised control of individual motor components^11–14^. The rate and pattern of SNr neuron firing *in vivo* is shaped by synaptic input. Canonically, the primary inputs to GABAergic SNr neurons originate from upstream basal ganglia nuclei, namely: the striatum, globus pallidus pars externa (GPe), and the subthalamic nucleus (STN)^15–17^. The SNr also receives input from structures outside the basal ganglia^18,19^. Compared to the intrinsic connectivity within the basal ganglia, little is known about the role that these external inputs play in modulating SNr activity.

Conventionally partitioned into primary and secondary subdivisions (herein abbreviated as M1 and M2, respectively), the rodent motor cortex is essential for motor learning and behavioural flexibility, as well as the coordination of skilled motor output^20–26^. Projection neurons of both M1 and M2 emit complex descending axons: individual neurons innervate several target structures through the basal ganglia, thalamus, midbrain, brainstem and spinal cord^26–31^. In rodents, anatomical evidence suggests that cortical axons also innervate the substantia nigra^14,26,27,30–33^. Retrograde transsynaptic tracing has identified both M1 and M2 as sources of input to the dopaminergic substantia nigra population^34^. However, whether the motor cortex forms functional synapses with the canonical GABAergic population of SNr neurons has not been explored.

In the present study, we sought to demonstrate functional synaptic transmission from the motor cortex to the GABAergic neurons of the SNr. We employed a virally targeted optogenetic approach, whereby we could selectively activate motor cortex efferents, and performed *ex vivo* whole cell recordings to assess postsynaptic responses in GABAergic SNr neurons. With the addition of pharmacological protocols, we establish the existence of a monosynaptic connection between glutamatergic motor cortex neurons and GABAergic SNr neurons, characterize the synaptic properties of this connection, and show that experimental activation of this pathway increases SNr firing rate *ex vivo*. To further understand the spatial distribution and projection patterns of cortical-recipient SNr neurons we employed a transsynaptic viral labelling approach. Our findings suggest that M1 and M2 target different subpopulations of SNr neurons. These results characterize functional cortical pathways that provide direct external regulation of basal ganglia output.

## Results

### Monosynaptic excitation of GABAergic SNr neurons by M1 projections

To gain an experimental handle on motor cortex efferents, we injected AAV2-CamKIIa-ChR2-eYFP unilaterally in M1 and obtained whole cell patch clamp recordings from SNr neurons in acute brain slices (Figure 1A, Bi). This approach offered expression of both the fluorescent reporter protein eYFP and the excitatory opsin channelrhodopsin (ChR2) selectively in M1 projection neurons, allowing us to simultaneously visualise and photostimulate efferent axons. Anatomically, we confirmed previous reports that M1 axons innervate the ipsilateral SNr, and recorded from SNr neurons located in the vicinity of labelled axons (Figure 1Bii-iv). The cell-type identity of recorded neurons was confirmed as either dopaminergic or GABAergic based on characteristic electrophysiological signatures (Figure 1C, S1A)^35,36^. Excitatory postsynaptic potentials (EPSPs) were evoked by widefield photostimulation, delivered as 20 Hz trains of 2 ms long light pulses (Figure 1D). Recordings were performed under constant bath application of the GABAA receptor antagonist gabazine (10 μM) to occlude concurrent synaptic activity stemming from either the striatum, GPe, or from neighbouring SNr neurons. In a subset of experiments (n = 9), EPSPs were abolished by bath application of the sodium channel antagonist Tetrodotoxin (TTX, 1 μM), confirming that the observed potentials were synaptic in nature. EPSPs could, however, be restored by subsequent bath application of the voltage-gated potassium channel antagonist 4-AP (100 μM), indicating that responses were monosynaptic in nature (Figure 1E)^37^. In a further subset of experiments (n = 8), EPSPs were robustly eliminated by bath application of glutamatergic receptor antagonists NBQX (10 μM) and D-APV (50 μM) (Figure 1F), confirming the glutamatergic nature of the synapses.

**Figure 1:**
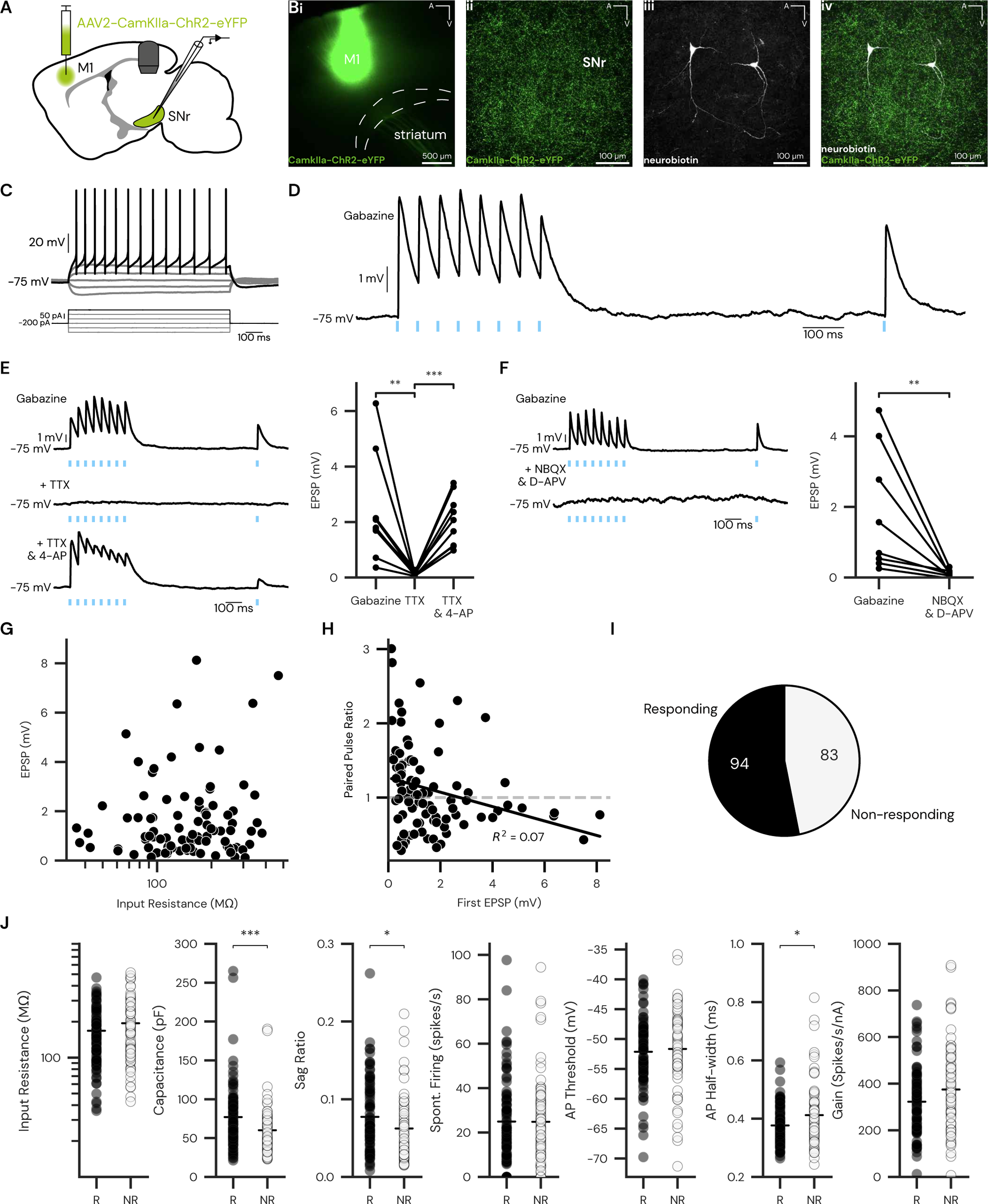
Photostimulation of M1 axon terminals evokes EPSPs in a subpopulation of GABAergic SNr neurons. **A**: Experimental setup. Channelrhodopsin (ChR2) expression was driven in projection neurons of M1 via injection of a viral vector (AAV2-CamkIIa-ChR2-eYFP). SNr neurons were recorded in acute brain slices via whole-cell patch clamp, in current clamp configuration. **B**: **i**. Representative widefield image of injection site in M1, in a parasagittal brain slice. **ii**. Confocal image of fluorescently-labelled axon terminals within the SNr **iii**. Addition of neurobiotin to the intracellular recording solution allowed for post-hoc visualization of neuron morphology. **iv**. Neuron morphology relative to fluorescently-labelled M1 axon terminals (A: anterior, V: ventral). **C**: GABAergic cell identity was assessed by characteristic responses to current pulses. **D**: Excitatory postsynaptic potentials (EPSPs) were recorded in response to trains of photostimulation (2 ms, 1 mW, stimulation onset indicated by blue marks) during bath application of the GABA_A_ antagonist gabazine (10µM). Hyperpolarizing current was injected to keep the baseline membrane potential at approximately −75 mV. **E**: Left: Representative response to photostimulation in the presence of gabazine (upper), gabazine and the sodium channel blocker TTX (1 µM) (middle), and gabazine, TTX and the voltage-gated potassium channel blocker 4-AP (100 μM) (lower). Right: summary of EPSP amplitudes recorded in the three conditions represented in the left panel (n = 9; ∗∗p < 0.01, ∗∗∗p < 0.001 two-tailed paired t-test). **F**: Left: photostimulation in the presence of gabazine (upper), and gabazine combined with glutamate receptor antagonists NBQX (10 µM) and D-APV (50 µM) (lower). Right: summary of EPSP amplitudes recorded in the three conditions represented in the left panel (n = 8; two-tailed paired t-test). **G**: Scatterplot of EPSP amplitudes and neuron input resistance from the recorded population. **H:** Scatterplot of first EPSP amplitude and paired pulse ratio. Solid black line indicates linear regression, horizontal dashed grey line indicates a paired pulse ratio of 1. **I**: Proportion of recorded neurons that responded to photostimulation; numbering represents total counts. **J**: Comparison of selected electrophysiological features between responding (R) and non-responding (NR) neurons (n = 94, N = 31; ∗p < 0.05, ∗∗∗p < 0.001, Mann-Whitney U). Mean values represented by horizontal bars.

From a holding potential of −75 mV, EPSPs presented with amplitudes ranging up to 8 mV. The recorded EPSP amplitudes were independent of the input resistance of the postsynaptic neurons (p = 0.11, r^2^ = 0.038; n = 94; N = 31; Figure 1G). This lack of correlation suggests that synaptic contacts are sparsely distributed across the dendritic arborisation^38,39^. SNr neurons possess long, passive dendrites, thought to play a role in integrating diverse signals^40,41^.

Photostimulation with a series of pulses at 20 Hz allows the engagement of short-term synaptic dynamics, while remaining within the kinetic activation range of the opsin (ChR2). The valence of short-term plasticity at corticofugal synapses is highly variable, even within the same postsynaptic cell type^42^. Similarly, when recording from SNr neurons, we observed both depression and facilitation (Figure 1H). A weak, inverse correlation was found between the measured paired pulse ratio and the amplitude of the first EPSP, with smaller amplitude synapses tending to exhibit facilitation (p = 0.008, r^2^ = 0.07, n = 94; N = 31).

The majority (53%) of GABAergic SNr neurons from which we recorded responded to photostimulation of cortical efferents (n = 94 of 177 neurons; Figure 1I). All neurons were subject to a battery of current injections to profile their intrinsic electrophysiological properties. Whereas most recorded neurons were GABAergic, a smaller number of dopaminergic neurons were recorded. Of this dopaminergic population, a minority responded to photostimulation of cortical terminals (n = 3 of 35 neurons; Figure S1B-D). Commonly reported sub- and suprathreshold electrophysiological properties were compared between responding (R) and non-responding neurons (NR). Responding neurons exhibited significantly higher membrane capacitance (R: 76.98 ± 4.25 pF, NR: 60.07 ± 3.18 pF; p = 0.0008), an increased sag ratio (R: 0.077 ± 0.005, NR: 0.062 ± 0.005; p = 0.023), and shorter action potential half-width (R: 0.377 ± 0.007 ms, NR: 0.412 ± 0.012 ms; p = 0.032) relative to non-responding neurons (n = 94 (R), n = 83 (NR); N = 31; Figure 1J). No significant differences were observed between the two groups in terms of input resistance, spontaneous firing rate and action potential threshold. Target-defined SNr subpopulations display distinct electrophysiological signatures^4^. Thus, the electrophysiological differences between responding and non-responding neurons may hint to a circuit motif whereby M1 selectively targets a specific set of SNr neurons.

Altogether, these results demonstrate that glutamatergic M1 projections form functional, monosynaptic connections with a population of GABAergic SNr neurons.

### M1 input to SNr neurons is mediated by AMPA receptors

The kinetics and short-term dynamics of postsynaptic responses to M1 input vary considerably between different postsynaptic cell types within the basal ganglia^42,43^. For a thorough characterisation of the M1-SNr synapse we recorded from SNr neurons in voltage clamp configuration while photostimulating M1 axon terminals (Figure 2A-B). From a holding potential of −70 mV, we recorded excitatory postsynaptic currents (EPSCs) with amplitudes spanning more than a 100-fold range (range: 6.7 - 680.4 pA; median: 58.1 pA; Figure 2C). EPSCs consistently presented with short onset latencies (as expected for monosynaptic connections), rise times and decay constants (Figure 2C). In voltage clamp configuration we detected EPSCs in 69% of neurons tested (n = 100 of 145; N = 20; Figure 2D). The increased proportion of respondent cells observed in voltage-clamp relative to current clamp configuration may reflect the fact that M1 terminals impinge upon the dendritic domain of SNr neurons. With a potassium-based intracellular solution, synaptic events arriving at distal dendrites may be less detectable due to dendritic filtering^44^. Our voltage-clamp intracellular solution was cesium-based, offering improved space clamping.

**Figure 2:**
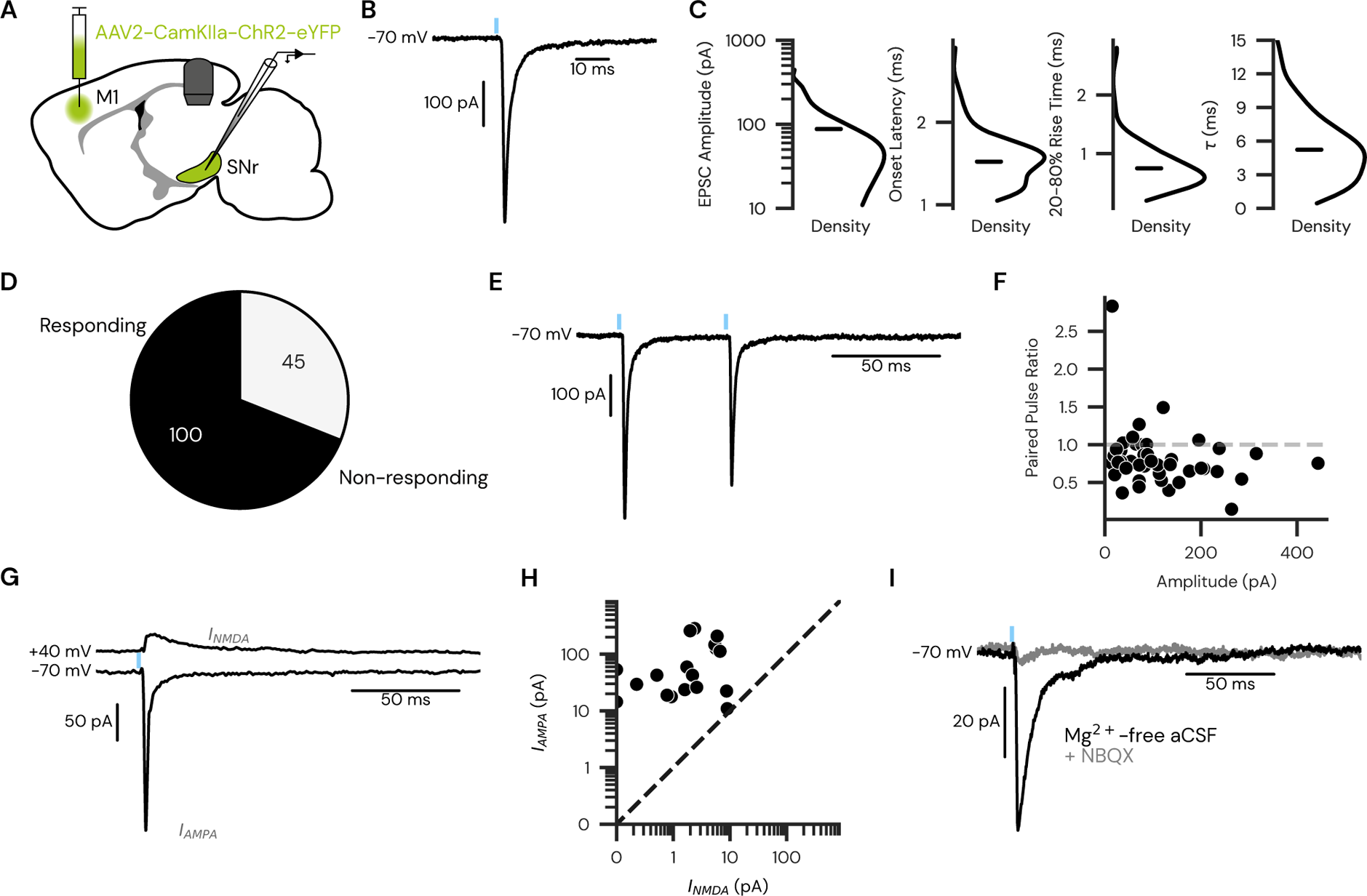
Characterisation of photoevoked EPSCs from M1 axon terminals. **A:** Experimental setup. ChR2 expression was driven in projection neurons of M1 via injection of a viral vector (AAV2-CamkIIa-ChR2-eYFP). SNr neurons were recorded in acute brain slices via whole-cell patch clamp, in voltage clamp configuration. **B:** Representative excitatory postsynaptic current (EPSC) recorded in response to photostimulation (2 ms, 1 mW, stimulation onset indicated by blue mark). **C:** Summary of EPSC amplitude and kinetics. Kernel density estimates for absolute EPSC amplitude, onset latency, 20-80% rise time and decay time constant (τ; n = 100). Mean values indicated by horizontal bars. **D:** Proportion of recorded neurons that responded to photostimulation in voltage clamp configuration; numbering represents total counts. **E:** In a subset of experiments, paired light pulses were delivered at a 50 ms interval (n = 44; 2 ms, 1 mW, stimulation onset indicated by blue mark). **F**: Scatterplot of first EPSC amplitude and paired pulse ratio for neurons tested with paired pulse protocol. Horizontal dashed grey line indicates a paired pulse ratio of 1 (n = 44). **G:** In a subset of experiments, EPSCs were recorded at both −70 and +40 mV to measure the relative contribution of AMPA and NMDA receptors. **H:** Scatterplot of absolute AMPA and NMDA currents recorded. Dashed black line indicates I_NMDA_ = I_AMPA_ (n = 18). **I:** In a further subset of experiments, EPSCs were recorded in a Mg^2+^-free solution before (black trace), and after bath application of the AMPA receptor antagonist NBQX (grey trace; n = 4).

In a subset of experiments, we again assessed the short-term plasticity of the M1-SNr synapse by measuring the paired pulse ratio in response to two consecutive light pulses (Figure 2E). We found a mean paired pulse ratio of 0.826 (n = 44), indicating a depressing synapse. Limited heterogeneity was observed: 8 of 44 neurons demonstrated facilitation. In comparing the paired pulse ratio to the amplitude of the first pulse we found no correlation (p = 0.11, r^2^ = 0.06; Figure 2F). These results are qualitatively different to what we observed in current clamp recordings (Figure 1H). Voltage clamp and current clamp recordings have previously been reported to yield different measures of short-term plasticity, an observation that has also been attributed to improved space clamping with voltage clamp intracellular solutions^45^.

Postsynaptic response to glutamatergic signalling is largely mediated by the expression of N-methyl-D-aspartate (NMDA) receptors and 〈-amino-3-hydroxy-5-methyl-4-isoxazolepropionic (AMPA) receptors^46^. To assess the relative contributions of these two receptor types at the M1-SNr synapse we separated the AMPA receptor-mediated and NMDA receptor-mediated EPSC components by photostimulating while holding the neuron at either negative or positive potentials (Figure 2G). We found a near complete absence of NMDA-mediated current. To further corroborate this result, we performed recordings in a Mg^2+^-free extracellular solution, thus removing the voltage-dependent Mg^2+^ gating of NMDA receptors^47,48^. Optically evoked EPSCs recorded in this Mg^2+^-free environment could be blocked by bath application of the AMPA receptor antagonist NBQX (Figure 3H, I). These results demonstrate that M1-SNr synaptic transmission is primarily AMPA receptor mediated, with fast kinetics.

**Figure 3:**
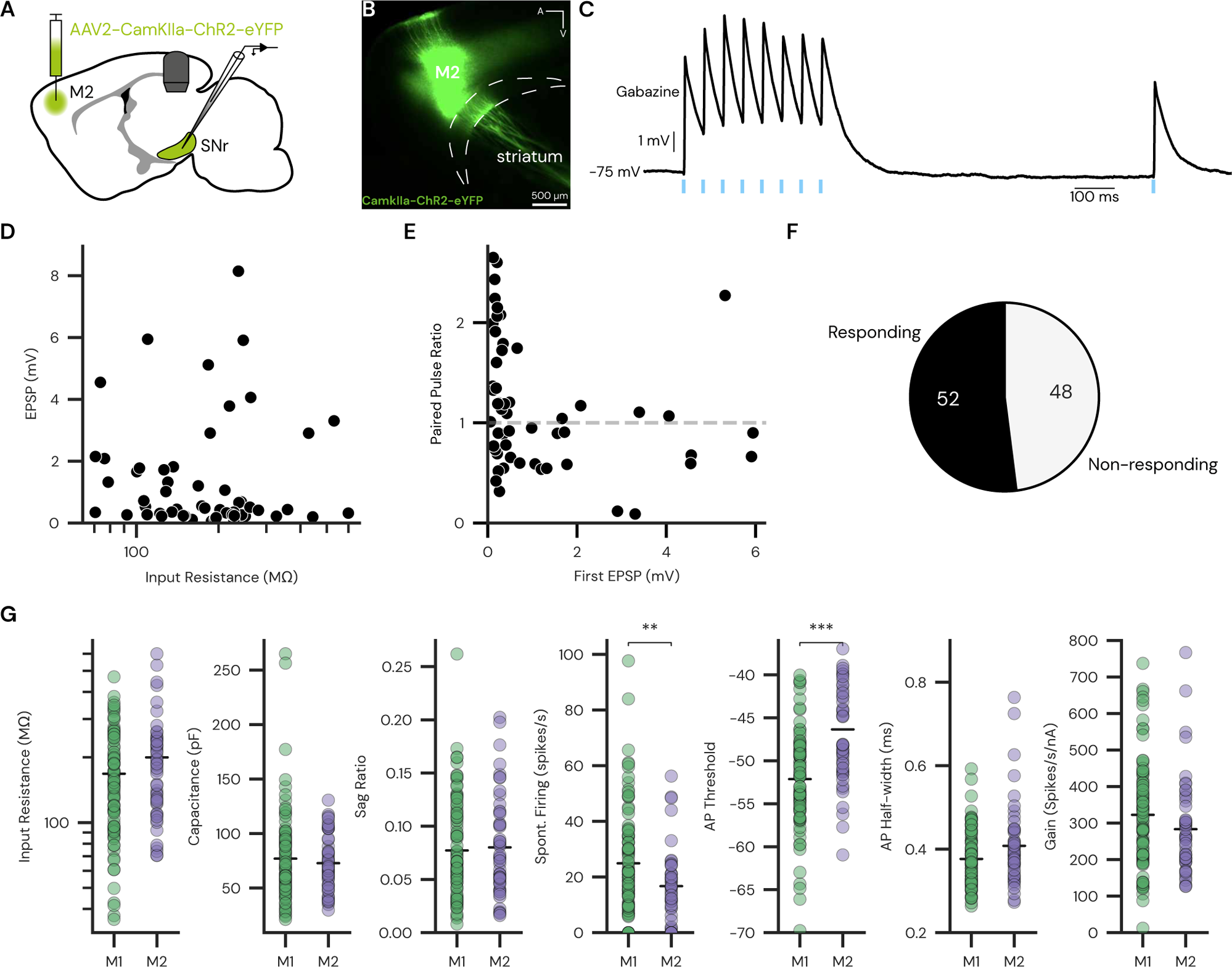
Photostimulation of M2 axon terminals evokes EPSPs in a subpopulation of GABAergic SNr neurons. **A**: Experimental setup. ChR2 expression was driven in projection neurons of M2 via injection of a viral vector (AAV2-CamkIIa-ChR2-eYFP; A: anterior, V: ventral). SNr neurons were recorded in acute brain slices via whole-cell patch clamp, in current clamp configuration. **B**: Representative widefield image of injection site in M2, in a parasagittal brain slice. **C**: EPSPs were recorded in response to trains of photostimulation (2 ms, 1 mW, stimulation onset indicated by blue marks) during bath application of the GABA_A_ antagonist gabazine (10 µM). Hyperpolarizing current was injected to keep the baseline membrane potential at approximately −75 mV. **D**: Scatterplot of EPSP amplitudes and neuron input resistance from the recorded population (n = 52, N = 20). **E:** Scatterplot of first EPSP amplitude and paired pulse ratio. Horizontal dashed grey line indicates a paired pulse ratio of 1. **F**: Proportion of recorded neurons that responded to photostimulation; numbering represents total counts. **G**: Comparison of selected electrophysiological features between M1-responding (M1) and M2-responding (M2) neurons (n = 94, 52; ∗p < 0.05, ∗∗p<0.01, ∗∗∗p < 0.001, Mann-Whitney U). Mean values represented by horizontal bars.

### A direct projection from M2 to SNr

The secondary motor cortex (M2) also has extensive direct projections to subcortical structures^49^. To assess whether M2 neurons target the GABAergic population of the SNr, we injected AAV2-CamKIIa-ChR2-eYFP unilaterally in M2, and recorded from SNr neurons, as previously detailed (Figure 3A, B). We recorded EPSPs in response to widefield photostimulation (Figure 3C). As with M1-SNr transmission, EPSP amplitude was not correlated to the input resistance of the postsynaptic neuron (p = 0.95, r^2^ < 0.0001; n = 52; N = 11 Figure 3D). M2-SNr EPSPs demonstrated a mix of facilitation and depression, with no correlation between the amplitude of the first EPSP in the train and the recorded paired pulse ratio (p = 0.22, r^2^ = 0.028; n = 52; N = 11; Figure 3E). 52% of neurons tested responded to M2 terminal photostimulation (n = 52 of 100 neurons; N = 11; Figure 3F).

Projections from M1 and M2 maintain topography as they innervate target structures in the subcortical motor system ^14,26,50^. As the intrinsic electrophysiological properties of SNr neurons also adhere to a topographical organisation^4,51^. We reasoned that M1 and M2 may target different populations within the SNr. We compared M1-responding to M2-responding neurons along common electrophysiological properties (Figure 3G). While no differences were observed in subthreshold properties, M1-responding cells displayed significantly higher spontaneous firing rate (M1: 24.9 ± 2.0 spikes/s, M2: 16.7 ± 2.6 spikes/s; p = 0.0038) and significantly lower action potential threshold (M1: −52.1 ± 0.6 mV, M2: −46.3 ± 0.9 mV; p < 0.0001; n = 94, N = 31 (M1); n = 52, N = 11 (M2); Figure 3G). Together, these results demonstrate that a population of SNr neurons receives synaptic input from M2, and that this population differs from the M1-recipient population.

### SNr neurons do not receive input from primary somatosensory cortex

To assess whether other cortical areas project to the SNr, we employed the same experimental strategy, but with the virus targeted to the primary somatosensory cortex (S1, Figure S2A). S1 projection neurons target the main input structure of the basal ganglia, the striatum, in a similar fashion to M1 projections^42^. However, S1 injections did not yield any labelled axons at the level of the SNr, and no recorded neurons responded to photostimulation (n = 27, N = 4; Figure S2B-C). Thus, cortical-SNr transmission appears to be unique to motor cortices.

### Photoactivation of motor cortex axon terminals increases SNr firing rate *ex vivo*

To assess whether cortical input is capable of shaping the firing patterns of SNr neurons, we employed the same viral approach to express ChR2 in either M1 or M2, and photostimulated ChR2-expressing axon terminals while recording the spontaneous activity of SNr neurons in cell-attached configuration (Figure 4A). In both M1 and M2 conditions, photoactivation at 20 Hz drove a transient increase in firing rate for the duration of the stimulus (n = 73; N = 21; and n = 20; N = 6, respectively; Figure 4A, B). The mean firing rate was significantly increased during the stimulation period relative to both pre- and post-stimulation periods (M1: pre: 19.4 ± 1.4 spikes/s, stimulation: 24.8 ± 1.5 spikes/s, post: 19.0 ± 1.4 spikes/s; pre - stim: p < 0.0001; stim - post: p < 0.0001; n = 73, N = 21; M2: pre: 10.1 ± 1.2 spikes/s, stimulation: 14.0 ± 1.5 spikes/s, post: 9.8 ± 1.3 spikes/s; pre - stim: p = 0.01; stim - post: p = 0.01; n = 20, N = 6; Figure 4C). Pre- and post-stimulation firing rates were not different in either condition (M1: p = 0.13; M2: p = 0.59). Similarly to what was observed in whole cell configuration, the baseline spontaneous firing rate of M1-recipient neurons was elevated relative to that of M2-recipient neurons (M1: 19.4 ± 1.4 spikes/s; M2: 10.1 ± 1.2 spikes/s; p = 0.001; Figure 4D). Thus, *ex vivo*, optogenetic activation of either M1 or M2 axon terminals can elevate the spiking activity of recipient SNr neurons.

**Figure 4:**
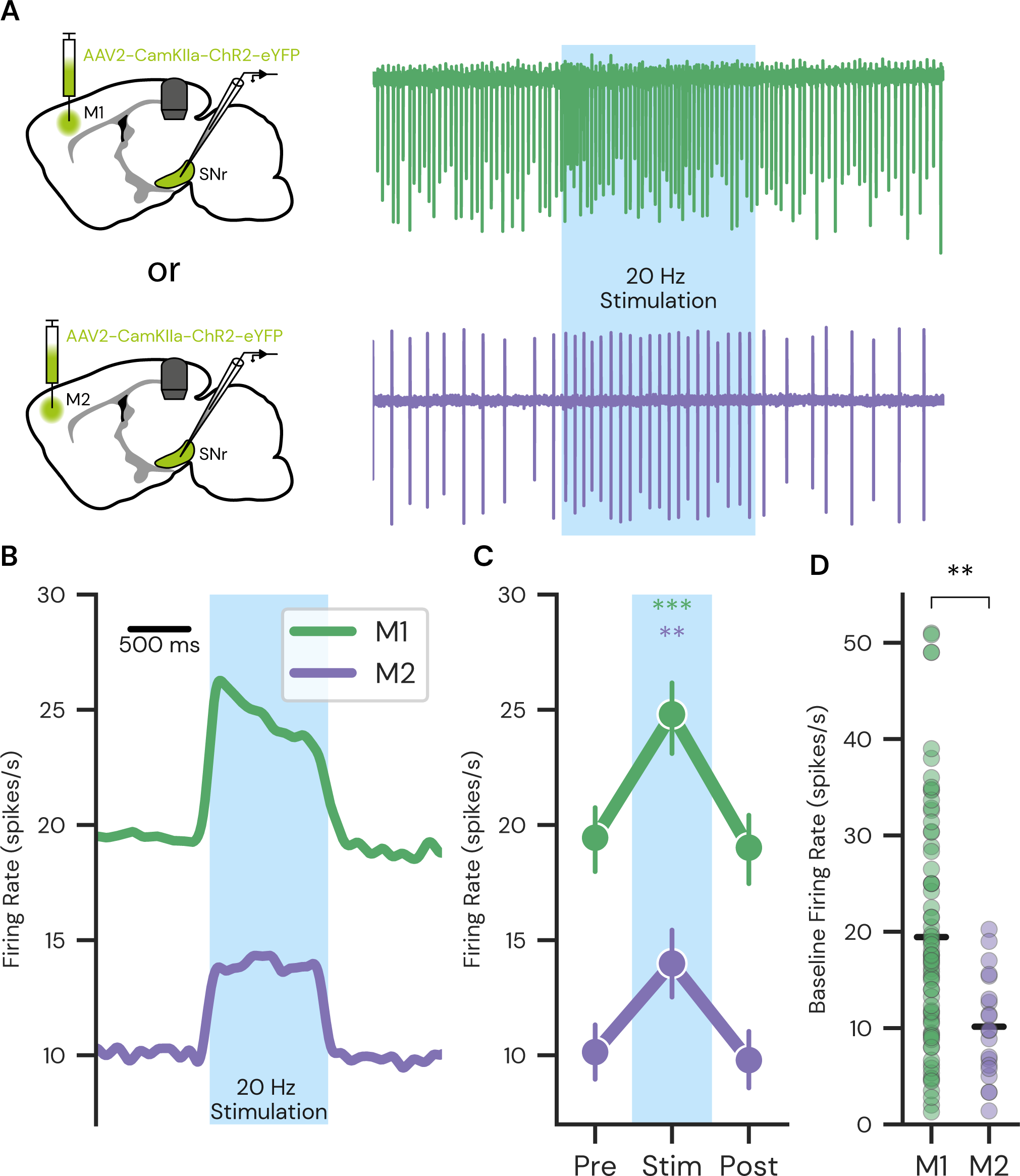
Photoactivation of cortical axon terminals increases intrinsic SNr firing. **A:** Experimental setup. ChR2 expression was driven in projection neurons of either M1 (upper left) or M2 (lower left) via injection of a viral vector (AAV2-CamkIIa-ChR2-eYFP) SNr neurons were recorded in cell-attached configuration and axon terminals were photostimulated at 20 Hz for one second (right). **B:** Mean firing rate response to photostimulation. Instantaneous firing rate was estimated per trial via kernel convolution (Gaussian kernel; σ = 50 ms) before averaging across trials, and across neurons (M1: green, n = 72; M2: magenta, n = 20). **C:** Binned firing rate during the stimulation period compared to pre- and post-stimulation (∗∗p < 0.01, ∗∗∗p < 0.001; two-tailed paired t-test,). Data presented as mean ± SEM. **D:** Comparison of pre-stimulation firing rate for M1- and M2-recipient neurons (∗∗p < 0.001; Mann-Whitney U).

### M1- and M2-recipient SNr neurons are topographically organised and have distinct projection patterns

To better understand the organisation of cortical input to SNr we employed an intersectional viral-labelling approach. We took advantage of the anterograde transsynaptic spread property of the AAV1 serotype to express Cre in neurons postsynaptic to either M1 or M2^52^. We combined this with a second AAV injection targeted to the ipsilateral SNr to selectively express a Cre-dependent fluorophore (GFP) in neurons receiving monosynaptic cortical input (Figure 5A). To map the spatial distribution of cortical-recipient somata, we registered individual sections to the Allen Mouse Brain common coordinate framework (version 3; Figure 5B). We verified the non-dopaminergic identity of labelled cells, via immuno-labelling for the dopaminergic marker tyrosine hydroxylase (TH): TH-negative, GFP-positive neurons were identified within the SNr, and their soma positions were recorded (Figure 5C, D). The spatial distribution of soma positions was quantified by kernel density estimates in the horizontal, and coronal planes (Figure 5E). As expected, M1- and M2-recipient neurons displayed a topography, with M1-recipient cells situated closer to the ventrolateral boundary of the nucleus (Figure 5E).

**Figure 5:**
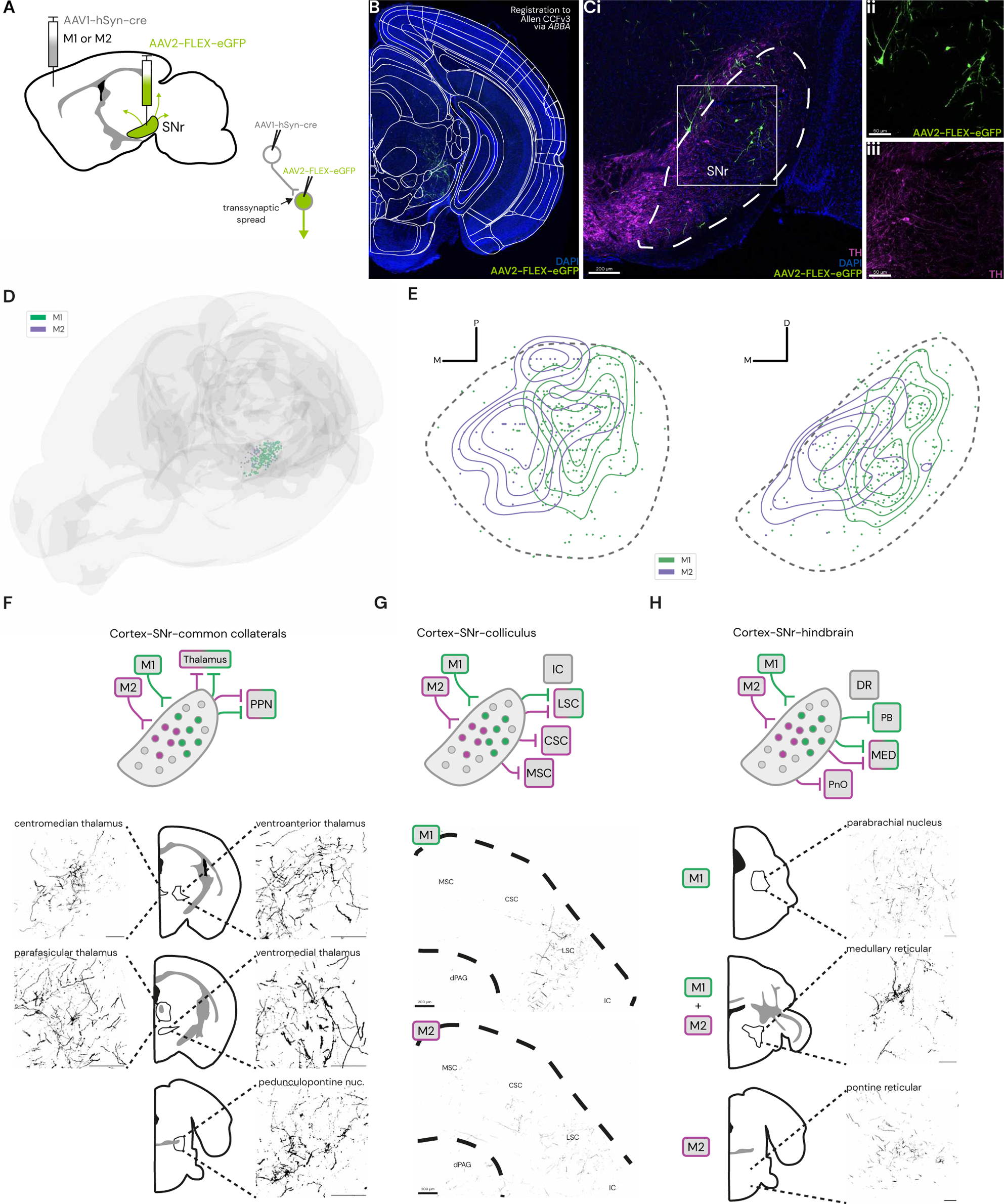
A transsynaptic viral strategy reveals topographical organisation of M1/M2-recipient SNr neurons. **A:** Experimental setup. AAV1-hSyn-Cre was injected in either M1 or M2. AAV2-FLEX-eGFP was injected in the ipsilateral SNr. Anterograde transsynaptic spread drives Cre expression in postsynaptic neurons, and fluorescent reporter protein expression in SNr neurons receiving monosynaptic input from the specific cortical region. **B:** Coronal brain sections containing the SNr were imaged and registered to the Allen Mouse Brain Common Coordinate Framework (v3) via *ABBA*. **C**: **i**. representative confocal image eGFP-expressing neurons and tyrosine hydroxylase (TH) reactivity in the SNr. **ii, iii**: magnified boxed region of **i**, showing isolated eGFP and TH channels. **D:** Distribution of eGFP-labelled somata for M1 (green) and M2 (magenta) injections in 3D (M1: N = 4, M2: N = 3). **E:** Distribution of eGFP-labelled somata projected on the coronal (left subpanel) or horizontal (right subpanel) plane. SNr boundaries are indicated by the dashed line. 2D-kernel density estimates are overlaid as contours with solid lines. **F:** eGFP-labelled axons observed in common collateral SNr targets in the thalamus and pedunculopontine nucleus. Representative images from M1-injected animals. Scale bars: 100 µm. **G:** eGFP-labelled axons innervation of the superior colliculus (upper: M1, lower: M2). **H:** eGFP-labelled axons observed in SNr targets in the pons/medulla. Scale bars: 100 µm (unless otherwise noted). cSC: *central superior colliculus,* dPAG: *dorsal periaqueductal grey,* DR: *dorsal raphe,* IC: *inferior colliculus,* lSC: *lateral superior colliculus,* mSC: *medial superior colliculus,* MED: *medullary reticular formation,* PB: *parabrachial nucleus,* PnO: *pontine reticular formation,* PPN: *pedunculopontine nucleus*.

We reasoned that the presence or absence of M1/M2 input may reflect differential downstream targeting of individual SNr neurons. Distinct SNr subpopulations project to unique sets of specialized targets in the brainstem, while sending collaterals to common targets in the thalamus and PPN^4,53^. We first verified that transsynaptically labelled neurons projected to common targets. Labelled axons were observed throughout motor-related thalamic nuclei, as well as in the PPN, in both M1- and M2-targeted experiments (Figure 5F).

We next focussed our analysis on specialized SNr targets in the colliculus and hindbrain. Primary SNr target structures of the colliculus, pontine and medulla receive input from largely mutually exclusive SNr populations, with minimal collateralisation between them^4^. For M1-injected brains, within the colliculus we observed fluorescently labeled axons exclusively in the lateral region of the superior colliculus (LSC; Figure 5G). The medial and central domains of the superior colliculus (CSC, MSC), as well as the inferior colliculus (IC), were void of fluorescence. In experiments where AAV1-Cre was injected to M2, however, labelled axons were observed broadly across the mediolateral extent of the SC (Figure 5G). No fluorescence was observed in the IC. Therefore, at the collicular level, while M1 input to SNr is specific to LSC-projecting SNr neurons, M2 projects to a wider range of SNr subpopulations. In the hindbrain, labelled axons from M1-recipient SNr neurons were found across the subdivisions of the parabrachial nucleus (Figure 5H). M1-recipient neurons were also found to target the medullary reticular formation - specifically concentrated around the parvicellular reticular nucleus (Figure 5H). M2-recipient neurons were also found to project through the hindbrain: labelled axons were seen passing through the pontine, with putative synapses in the pontine reticular formation (Figure 5H). Labelled fibers were also seen extending to the level of the rostral medulla. Altogether, these findings indicate that M1 and M2 project to largely separate SNr subpopulations.

## Discussion

To understand cortical control of downstream motor centers requires a thorough account of the descending connectivity of motor cortex. Similarly, to fully understand how SNr activity is regulated, a complete description of SNr inputs is necessary. In this study we have confirmed a direct, functional connection between these two brain regions, by combining virally targeted optogenetics and *ex vivo* electrophysiology. Our results show that subpopulations of GABAergic SNr neurons receive monosynaptic, glutamatergic input from the primary and secondary motor cortices and demonstrate that this input is capable of shaping SNr neuron firing patterns. Using a transsynaptic-labelling approach we identified downstream targets of M1-recipient and M2-recipient SNr neurons.

Our anterograde transsynaptic approach identified a disynaptic M1-SNr-LSC pathway. This is corroborated by our finding that SNr neurons responding to photostimulation of M1 terminals have an inherently higher membrane capacitance than non-responding neurons. In a study combining retrograde labelling and *ex vivo* electrophysiology McElvain et al. reported that LSC-projecting SNr neurons have significantly greater membrane capacitances than the SNr population as a whole^4^. Thus, our anatomical findings agree with our electrophysiological data. M2-recipient SNr neurons were found to project broadly across the SC. Using an identical viral-tracing strategy, Lee et al. demonstrated a similar topography in SNr-SC projections, with the SNr neurons that receive input from the ventrolateral striatum targeting the LSC, while SNr neurons receiving input from dorsal regions of the striatum project broadly across the SC^11^. The colliculus also receives monosynaptic input from the cortex.

Corticocollicular projections maintain a strict topography, whereby M1 neurons synapse exclusively in the lateral SC^4,54^. M2-SC projections are broader, targeting the lateral, central and medial SC^54^. Therefore, the monosynaptic corticocollicular and disynaptic cortico-nigro-collicular pathways remain in register, converging in the same region of the SC.

Information flow through the basal ganglia circuit has traditionally been thought of in terms of competing pathways ^55–59^. The disynaptic Hyperdirect (cortex-STN-SNr) pathway provides excitatory input to SNr neurons via glutamatergic neurons in the STN^59,60^. This excitatory input drives increased SNr firing and ultimately elicits increased inhibition of both thalamic nuclei and downstream motor targets. The Hyperdirect pathway therefore provides a route for a cortical command to either suppress competing motor programs through inhibition of motor centres or interrupt ongoing movement via the SNr-thalamus-cortex loop^60,61^. The existence of a monosynaptic corticonigral connection may serve as a reinforcement of the Hyperdirect pathway. Intersectional retrograde labelling studies in mice have demonstrated that M1 neurons innervating the SNr also innervate the STN^14,26^. Whether individual SNr neurons receive concurrent excitation through both subthalamonigral and corticonigral inputs has not been shown, although the STN appears to target all neurons of SNr^51^. Although STN input is a powerful driver of SNr neurons, STN neurons are also subject to competitive inhibition from the prototypical neurons of the GPe. A functional advantage of having a monosynaptic pathway for excitation, may be that a cortical command could reach the SNr unperturbed, providing robust excitatory drive independently of concurrent activity in the other nodes of the basal ganglia circuit.

Cortical projection neurons also target GABAergic neurons of the striatum and GPe^28,42,62^. Cellular reconstructions offer some evidence to suggest that individual M1 neurons projecting to the striatum and those projecting to the SNr are largely mutually exclusive populations^31^. GPe-projecting cortical neurons showed diverse collateralization patterns, with some neurons collateralizing in SNr. An electrophysiological study in rats found that, within the GPe, input from motor cortices was biased towards the striatum-projecting ‘arkypallidal’ neurons, which are not known to project to SNr^28^. In contrast, multiple studies in mice have reported cortical input to both major GPe cell classes^31,62^. Under our experimental conditions, axons projecting to the SNr from both the striatum and the GPe were typically severed by the slicing process. In addition, GABAergic synapses were blocked by constant bath application of gabazine, as previously mentioned.However, these experimental constraints are not representative of normal physiological activity. Both M1 and M2 contain a diversity of unique projection neuron populations that are activated dynamically in behavioural contexts^63–65^. Non-specific experimental activation of M1 or M2 in the intact brain would conceivably excite all basal ganglia structures. Electrical stimulation of the rodent motor cortex *in vivo* generates multiphasic responses in the SNr^66,67^. As we have shown that corticonigral input is monosynaptic, the time delay between cortical activation and SNr response would be minimal. Thus, the early excitation phase commonly attributed to the Hyperdirect pathway likely also encompasses monosynaptic input from the corticonigral pathway. This may explain the persistence of an early excitation response in SNr neurons of mice following selective ablation of the Hyperdirect pathway^60^.

Alternatively, monosynaptic cortical input to the SNr could be reflective of a broader circuit motif for cortical orchestration of the brain’s motor system. By simultaneously activating multiple downstream structures, the motor cortex ensures that motor commands are broadcast to all relevant targets together^68^. The motor cortex exhibits increased activity in advance of motor execution^69,70^. This type of command could prime SNr neurons to be in a specific activity state in anticipation of forthcoming signals from, for example, the striatum. By reinforcing an SNr neuron’s firing rate, cortical commands would ensure that the neuron would be in a position to respond to the following striatal inhibition.

Input arriving directly from the cortex would provide higher temporal fidelity than signals that must first be processed in upstream basal ganglia nuclei. The ability to perform complex movements with high temporal precision may depend on this unhindered communication between the cortex and subcortical targets. The cortical neurons that innervate SNr were shown to be located in motor regions relating to the control of the forelimbs and mouth^14,26^. This is complemented by the higher density of cortical axon innervation observed in the lateral regions of the SNr. These studies noted an absence of cortical axon terminals in the most medial regions of SNr. Our findings further confirm that only neurons in the lateral portion of SNr receive monosynaptic input from the motor cortex. The medial SNr has been implicated in associative functions and behavioural state transitions, as well as gross locomotion^18,71^. Medially located neurons fire at lower rates than their lateral counterparts, and project to different targets in the brainstem^4^. Relative to their lateral counterparts, medial SNr neurons receive increased input from hypothalamic and state-related midbrain regions, and are involved in the processing of homeostatic signals with slow time resolution^18^. Signals with high temporal precision -i.e., monosynaptic cortical input – would logically be targeted towards the SNr neurons participating in motor circuits. While we report that the majority of SNr cells assessed responded to photostimulation of M1/M2 axon terminals, this number is biased by our ability to visualise fluorescent terminals in the slice before performing recordings and therefore does not reflect a uniform sampling of the SNr population as a whole.

As mentioned, all our recordings were conducted in the presence of the GABAA receptor antagonist gabazine. This prevented tonic GABAergic input from shunting or otherwise occluding optically evoked synaptic responses. However, under physiological conditions, simultaneous excitation of a number of SNr neurons by cortical input could lead to enhanced lateral GABAergic signalling between SNr neurons. The combined effect of cortical excitation and local inhibition remains unclear. Furthermore, the number of SNr neurons targeted by a single cortical axon is unknown. The functional relevance of local inhibition in this context may depend on the size of the SNr neuron population simultaneously recruited by cortical input. However, local connectivity within the SNr is sparse, and may reflect competition between parallel modules^72^. Whether connections between SNr neurons that share a common downstream target exist is unknown.

All experiments in the current study were conducted *ex vivo*. While we were able to demonstrate that optogenetic activation of cortical input is capable of significantly increasing SNr neuron firing rate, the behavioural implications of this connection remain unclear. Future work could show the functional relevance of this connection by selectively interfering with cortical signalling at the level of the SNr in awake, behaving animals.

## Methods

### Materials availability

This study did not generate new unique reagents.

### Experimental model and subject details

All experiments were performed with approval of the local ethical board, Stockholm Norra Djurförsöksetiska Nämnd, under an ethical permit to Gilad Silberberg (N2022-2020). Both male and female mice were used in this study. Wild type mice (C57BL/6J, stock #000664, The Jackson Laboratory) were used. Mice were housed in groups of up to four and maintained on a 12-h light cycle with *ad libitum* access to standard food and water.

### Method details

#### Virus injections

Mice, aged between six and ten weeks, were anaesthetised with isoflurane (VM Pharma AB, Sweden), and placed in a stereotaxic frame (Stoelting, U.S.A). Craniotomies were drilled with the following coordinates relative to bregma: M1 (AP: 1.5 mm, ML: −1.75 mm, DV: −0.8 mm), M2 (AP: 2.3 mm, ML: −1.00 mm, DV: −0.9 mm), SNr (AP: −3.4 mm, ML: −1.5 mm, DV: −4.5 mm). For electrophysiological experiments 0.4 µL of AAV2-CaMKIIa-hChR2(H134R)-eYFP (Addgene #26969; Penn Vector Core, USA) was injected into either M1 or M2. For transsynaptic labelling experiments 0.4 µL of AAV1-hSyn-Cre (Addgene #105553; Penn Vector Core, USA) was injected into either M1 or M2, and 0.1 µL of AAV2-pCAG-FLEX-eGFP (Addgene #51502; titer: 7×10¹² vg/mL) was injected into the SNr during the same surgery. For all injections, viruses were injected using a micropipette, at a rate of 0.1 µL/min (Quintessential Stereotaxic Injector, Stoelting, U.S.A.). The pipette was held in place for at least 5 min before being slowly retracted from the brain. Buprenorphine (Indivior Europe Limited; Apoteket, Sweden) was administered perioperatively (0.1 mg/kg).

#### Brain slice preparation

Mice were sacrificed at least three weeks following virus injections. Mice were anaesthetised with isoflurane (VM Pharma AB, Sweden) and decapitated. Brains were removed while immersed in ice-cold cutting solution containing the following (in mM): 2.5 KCl, 1.25 NaH_2_PO_4_, 0.5 CaCl_2_, 7.5 MgCl_2_, 10 glucose, 25 NaHCO_3_, 205 sucrose. Either coronal or parasagittal slices were cut at a thickness of 250 µm with a Leica VT1200S Vibratome (Leica Microsystems GmbH, Germany). Slices were incubated for 30 minutes at 35°C in a chamber filled with artificial cerebrospinal fluid (aCSF) saturated with 95% oxygen and 5% carbon dioxide. The aCSF contained the following (in mM): 125 NaCl, 25 glucose, 25 NaHCO_3_, 2.5 KCl, 2 CaCl_2_, 1.25 NaH_2_PO_4_, 1 MgCl_2_. Before recording, slices were kept in aCSF at room temperature for at least 45 minutes.

#### *Ex vivo* recordings

Whole-cell patch clamp recordings were obtained in oxygenated aCSF, which was continuously perfused throughout the experiment and maintained at 35°C with a temperature control unit (Luigs and Neumann GmbH, Germany). Neurons were visualised using infrared differential interference contrast (IR-DIC) microscopy on a BX51WI (Olympus, Japan) upright microscope, with a 40× long-working-distance immersion objective and a digital camera (Hamamatsu Photonics, Japan). Borosilicate glass pipettes were pulled using a P1000 micropipette puller (Sutter Instrument, U.S.A.) to a resistance of 6–8 MΩ and filled with an intracellular solution containing (in mM): 130 K-gluconate, 5 KCl, 10 HEPES, 4 Mg-ATP, 0.3 GTP, 10 Na_2_-phosphocreatine. For post-hoc localisation and morphological analysis, 0.2% neurobiotin (Vector Laboratories, U.S.A.) was added to the intracellular solution in a subset of experiments. Liquid junction potential (measured at approximately 11 mV) was not corrected for. For experiments conducted in voltage clamp configuration, a cesium-based internal solution was used instead, and contained (in mM): 10 CsCl, 110 CsMeSO_3_, 10 HEPES, 10 Na_2_-Phosphocreatine, 4 ATP-Mg, 0.3 GTP-Na, 10 TEA-Cl, 1 QX-314-Cl.

SNr neuron identity was confirmed by characteristic electrophysiology (. Up to three cells were recorded simultaneously. Recordings were performed with MultiClamp 700B amplifiers (Molecular Devices, U.S.A.), filtered at either 4 kHz (voltage clamp) or 10 kHz (current clamp), and digitised (10–20 kHz) via an ITC-18 (HEKA Elektronik, U.S.A.) acquisition board. Recordings were acquired with IGOR Pro 6.37 (WaveMetrics, U.S.A.).

Intrinsic neuronal properties were extracted in whole cell current clamp configuration using a standardised series of hyperpolarizing and depolarizing current injections. A baseline hyperpolarizing current was injected to maintain resting membrane potentials at approximately −75 mV. To activate ChR2, 473 nm light was transmitted to the slice via the objective (CoolLED Limited, U.K.). Trains of light pulses (2 ms duration, 8 pulses) were delivered at 10, 20 or 40 Hz. For voltage clamp experiments, a single light pulse was delivered. In all conditions responses were averaged from a minimum of 5 sweeps, with a 10 second interval between consecutive sweeps. In a subset of experiments, light stimulation was delivered at 20 Hz for 1 second while holding the cell in cell-attached configuration, before moving to whole-cell configuration. Pharmacological agents were bath applied to the slice, via the perfusate. These included tetrodotoxin (1 μM; Tocris, U.K.) and 4-aminopyridine (100 μM; Tocris, U.K.) to isolate monosynaptic responses, NBQX (10 μM; Tocris, U.K.) and D-APV (50 μM, Tocris), to selectively block glutamatergic signalling, and gabazine (SR-95531; 10 μM; Sigma-Aldrich, U.K.) to selectively block GABA_A_ receptor signalling. For morphological analysis, 0.2% neurobiotin (Vector Laboratories, U.S.A.) was added to the intracellular solution in a subset of experiments. The recorded slices were fixed overnight in 4% (w/v) paraformaldehyde (PFA) with picric acid, washed in 0.01 M phosphate buffered saline (PBS, pH 7.3), and incubated for 48 to 72 hours at 4°C with Cy5-conjugated streptavidin (1:1000, Jackson ImmunoResearch Laboratories) in 0.01 M PBS containing 0.6% Triton X-100. Following three washes in PBS, slices were counterstained with DAPI (1:5000; Sigma-Aldrich, Cat: D9542) and mounted on Superfrost Plus microscope slides (Thermo Fisher Scientific, USA.) with ProLong Antifade mounting agent (Thermo Fisher Scientific, USA). Confocal z-stacks were taken on either a Zeiss LSM 700 or LSM 800 microscope, running ZEN Blue (Zeiss, Germany).

#### Histology

For viral tracing experiments, animals were anesthetized and transcardially perfused with 4% PFA in 0.01 M PBS, at least three weeks after virus injections. Brains were removed and post-fixed overnight in PFA at 4°C. Brains were then transferred to a sucrose/PBS solution (30% w/v) for 18-24 hours. Serial 50 µm coronal or sagittal sections were cut on a cryostat. To discriminate between GABAergic and dopaminergic neurons, certain sections were stained for tyrosine hydroxylase (TH) immunoreactivity. Sections were incubated in blocking solution (5% normal donkey serum and 0.3% Triton X-100 in PBS) for 30 minutes and then incubated in rabbit anti-TH polyclonal antibody (Millipore, Cat: AB152) diluted 1:1000 in 0.3% Triton X-100 in PBS, at 4 °C, overnight. Sections were rinsed in PBS and incubated in either Cy3-conjugated goat, or Cy5-conjugated donkey, anti-rabbit polyclonal secondary antibody (The Jackson Laboratory, Cat: 111-165-003) for two hours at room temperature. Sections were stained with DAPI (1:5000; Sigma-Aldrich, Cat: D9542) and mounted with ProLong Antifade mounting agent (Thermo Fisher Scientific, USA). Confocal tile scans were taken on a LSM 800 microscope, running *ZEN Blue* (Zeiss, Germany). Complimentary widefield images were taken with an Olympus XM10 digital camera mounted on an Olympus BX51 fluorescence microscope (both Olympus Sverige AB, Stockholm, Sweden). Image brightness and contrast were adjusted in *ImageJ*^73^. Sections were registered to the Allen Mouse Brain Common Coordinate Framework (Version 3) using the *ImageJ* macro *ABBA*^74^.

#### Quantification and Statistical Analysis

Neuron distribution was quantified using *QuPath*^75^ and visualised with *Blender 4.0*. All other analysis was performed using custom written Python scripts (Python 3.9 and 3.11), using open-source packages from *eFEL, Elephant* and *NEO*^76–78^. Results are presented in the text as mean ± SEM. Statistical significance is defined as ∗ p < 0.05, ∗∗ p < 0.01, and ∗∗∗ p < 0.001.

## Data and Code Availability

All acquired data and generated code are available upon request.

## Acknowledgments

We thank Elin Dahlberg and Kristoffer Tenebro Berglund for technical assistance and animal colony management, and Dr. Brita Robertson for comments on the manuscript. This work was supported by the Swedish Medical Research Council (VR-M-2019-01854, VR-M-2019-01254, VR-M-2023-02304, VR-M-2021-01995), EU/FP7 Moving Beyond grant ITN-No-316639, under grant agreement no. 604102 (HBP), EU/Horizon 2020, no. 945539 (HBP SGA3), the European School of Network Neuroscience (MSCA-ITN-ETN H2020-860563), Knut & Alice Wallenberg Stiftelse (KAW 2017.0273), the Swedish Brain Foundation (Hjärnfonden, FO2023-0230), and grants from Karolinska Institutet.

## Author Contributions

W.S.T., S.G. and G.S. conceived the experiments. W.S.T. performed experiments and analysis. W.S.T., S.G. and G.S. wrote the manuscript.

