## Supplementary Figures for "Motor cortex directly excites the output nucleus of the basal ganglia, the substantia nigra pars reticulata"

Supplementary material consisting of 2 figures.

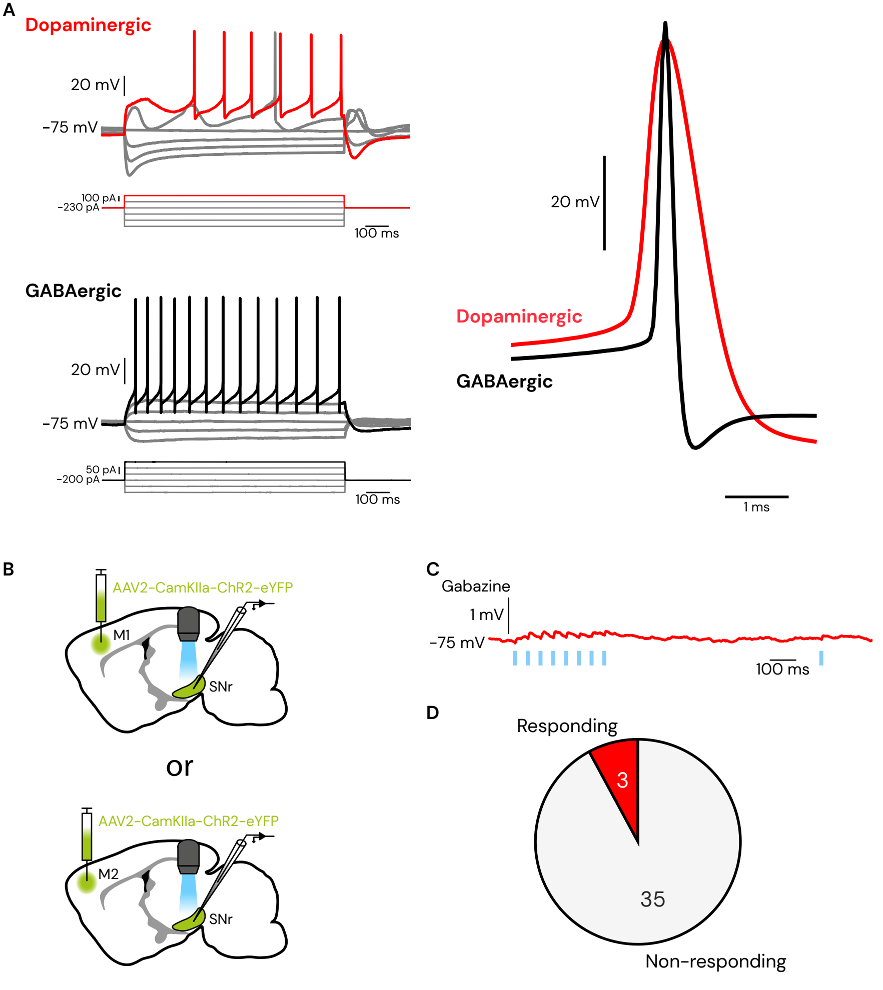

A
Dopaminergic
20 mV
-75 mV
100 pA
-230 pA
100 ms
GABAergic
20 mV
-75 mV
50 pA
-200 pA
100 ms
20 mV
Dopaminergic
GABAergic
1 ms
B
AAV2-CamKIIa-ChR2-eYFP
M1
SNr
or
AAV2-CamKIIa-ChR2-eYFP
M2
SNr
C
Gabazine
1 mV
-75 mV
100 ms
D
Responding
3
35
Non-responding

**Supplementary Figure 1: Dopaminergic neurons within the SNr are electrophysiologically distinguishable from GABAergic neurons**

**A**: Electrophysiological characterisation of dopaminergic and GABAergic neurons in the substantia nigra. Left: representative responses to current steps for a dopaminergic (upper) and GABAergic (lower) neuron. Right: representative single action potentials for a dopaminergic (red) and GABAergic (black) neuron. **B**: Experimental setup. ChR2 expression was driven in projection neurons of either M1 or M2 via injection of a viral vector (AAV2-CamkIIa-ChR2-eYFP). SNr neurons were recorded in acute brain slices via whole-cell patch clamp, in current clamp configuration. **C**: EPSPs were recorded in response to trains of photostimulation (2 ms, 1 mW, stimulation onset indicated by blue marks) during bath application of the $GABA_A$ antagonist gabazine (10 μM). Hyperpolarizing current was injected to keep the baseline membrane potential at approximately -75 mV. **D**: Proportion of recorded dopaminergic neurons that responded to photostimulation; numbering represents total counts; counts are pooled between M1 and M2 injections.

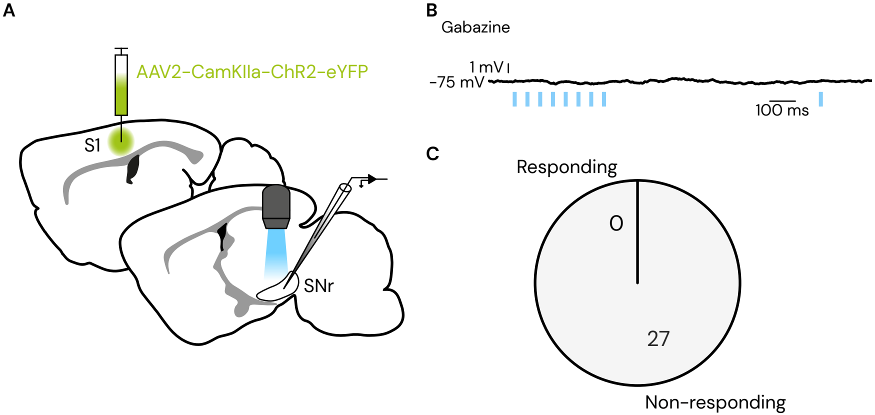
A
AAV2-CamKIIa-ChR2-eYFP
S1
SNr
B
Gabazine
1 mV
-75 mV
100 ms
C
Responding
0
27
Non-responding

**Supplementary Figure 2: Photostimulation of S1 axon terminals does not produce excitation of SNr neurons**

**A**: Experimental setup. ChR2 expression was driven in projection neurons of S1 via injection of a viral vector (AAV2-CamkIIa-ChR2-eYFP). SNr neurons were recorded in acute brain slices via whole-cell patch clamp, in current clamp configuration. **B**: No EPSPs were observed in response to trains of photostimulation (2 ms, 1 mW, stimulation onset indicated by blue marks) during bath application of the $GABA_A$ antagonist gabazine (10 μM). Hyperpolarizing current was injected to keep the baseline membrane potential at approximately -75 mV. **C**: Proportion of recorded neurons that responded to photostimulation; numbering represents total counts.
